## Supplementary Tables for "The neural dynamics of positive and negative expectations of pain"

### Supplementary Material

### Supplementary Table 1

*Peak coordinates and statistics of regions that showed common effects of positive and negative expectations compared to control in the anticipation phase*

| | | MNI Coordinates | | | | |
|---|---|---|---|---|---|---|
| **Region** | **Hemi.** | **X** | **Y** | **Z** | ***t*** | $p_{FWE}$ |
| *Expectation > Neutral Expectation* | | | | | | |
| Angular Gyrus | R | 56 | -54 | 40 | 5.92 | <.001 |
| Superior Frontal Gyrus | R | 8 | 44 | 36 | 5.57 | .002 |
| | L | -4 | 30 | 62 | 4.88 | .049 |
| Insular Cortex | L | -28 | 22 | -6 | 5.33 | .006 |
| Paracingulate Gyrus | R | 6 | 46 | 12 | 5.02 | .027 |
| vmPFC | R | 14 | 56 | -14 | 5.00 | .029 |
| Anterior Cingulate Cortex | R | 4 | 42 | 12 | 4.89 | .047 |
| | L | -2 | 40 | -4 | 3.96 | .025† |
| DLPFC | R | 40 | 24 | 36 | 4.44 | .004† |
| | L | -32 | 18 | 36 | 3.93 | .026† |
| Thalamus | L | -6 | -12 | 4 | 3.91 | .038† |
| *Neutral Expectation > Expectation* | | | | | | |
| Superior Parietal Lobule, Postcentral Gyrus | R | 34 | -36 | 44 | 5.62 | .001 |
| DLPFC | R | 30 | 2 | 62 | 3.74 | .048 |

*Note.* Coordinates are in MNI space. DLPFC = Dorsolateral Prefrontal Cortex. vmPFC = ventromedial prefrontal cortex. † small-volume corrected.

**Supplementary Table 2**

*Peak coordinates and statistics of regions that showed differential activation for expectation compared to neutral expectation in the pain phase*

| | | MNI Coordinates | | | | |
|---|---|---|---|---|---|---|
| **Region** | **Hemi.** | **X** | **Y** | **Z** | ***t*** | ***$p_{FWE}$*** |
| *Expectation > Neutral Expectation* | | | | | | |
| Brain Stem | L | -14 | -28 | -30 | 4.14 | .026† |
| Thalamus | R | 4 | -4 | -2 | 4.05 | .023† |
| *Neutral Expectation > Expectation* | | | | | | |
| Precentral Gyrus | R | 24 | -6 | 44 | 5.16 | .014 |
| | L | -60 | -2 | 24 | 5.05 | .023 |
| Amygdala | L | -18 | 0 | -20 | 4.28 | .003† |
| Hippocampus | R | 32 | -22 | -14 | 4.19 | .009† |
| Insular Cortex | R | 46 | -10 | 16 | 4.01 | .041† |
| DLPFC | R | 42 | 10 | 24 | 3.82 | .037† |

*Note.* Coordinates are in MNI space. DLPFC = Dorsolateral Prefrontal Cortex. † small-volume corrected.

**Supplementary Table 3**

*Peak coordinates and statistics of regions that showed differential activation between placebo and nocebo in the anticipation phase*

| | | MNI Coordinates | | | | |
|---|---|---|---|---|---|---|
| **Region** | **Hemi.** | **X** | **Y** | **Z** | ***t*** | **$p_{FWE}$** |
| *Placebo > Nocebo* | | | | | | |
| Lingual Gyrus | L | -14 | -76 | -6 | 11.34 | <.001 |
| Occipital Fusiform Gyrus | L | -20 | -68 | -12 | 9.21 | <.001 |
| Precuneus | L | -2 | -62 | 16 | 5.42 | .004 |
| Supracalcarine Cortex | R | 14 | -62 | 16 | 5.05 | .024 |
| Superior LOC | L | -16 | -84 | 24 | 4.96 | .034 |
| Amygdala | R | 22 | 2 | -22 | 3.62 | .031† |
| *Nocebo > Placebo* | | | | | | |
| Lingual Gyrus | R | 10 | -80 | -6 | 13.27 | <.001 |
| LOC | R | 28 | -78 | 20 | 5.46 | .003 |
| Cerebellum VI | R | 34 | -62 | -20 | 5.09 | .019 |

*Note.* Coordinates are in MNI space. LOC = Lateral Occipital Cortex. † small-volume corrected.

**Supplementary Table 4**

*Peak coordinates and statistics of regions that showed larger activation for positive expectations compared to negative expectations in the pain phase*

| Region | Hemi. | MNI Coordinates X | MNI Coordinates Y | MNI Coordinates Z | *t* | $p_{FWE}$ |
|---|---|---|---|---|---|---|
| *Placebo > Nocebo* | | | | | | |
| Anterior SMG | L | -36 | -40 | 36 | 6.53 | <.001 |
| DLPFC | R | 40 | 14 | 46 | 4.80 | .001† |
| | L | -22 | 8 | 44 | 6.29 | <.001† |
| Superior Parietal Lobule | R | 30 | -64 | 46 | 6.17 | <.001 |
| | L | -30 | -60 | 44 | 5.16 | .014 |
| Middle Frontal Gyrus | L | -40 | 12 | 36 | 6.14 | <.001 |
| Angular Gyrus | R | 44 | -50 | 38 | 6.14 | <.001 |
| vmPFC | R | 42 | 46 | 10 | 5.72 | .001 |
| | L | -42 | 50 | 2 | 6.00 | <.001 |
| Precuneus | R | 4 | -66 | 48 | 5.49 | .003 |
| | L | -20 | -66 | 56 | 5.70 | .001 |
| Posterior SMG | R | 42 | -36 | 48 | 5.24 | .010 |
| Superior Frontal Gyrus | L | -24 | 6 | 56 | 5.19 | .012 |
| Middle Temporal Gyrus | R | 60 | -50 | -10 | 5.17 | .014 |
| Brain Stem | R | 4 | -42 | -38 | 4.06 | .035† |
| | L | -2 | -42 | -34 | 5.16 | .014 |
| Inferior Frontal Gyrus | R | 44 | 12 | 18 | 4.89 | .047 |
| | L | -44 | 34 | 14 | 4.97 | .033 |
| Anterior STG | L | -60 | -10 | 2 | 4.88 | .048 |
| Hippocampus | R | 28 | -16 | -18 | 4.81 | .001† |
| | L | -24 | -18 | -16 | 3.82 | .032† |
| Central Operculum | L | -60 | -10 | 8 | 4.19 | .021† |
| Insula | L | -40 | -2 | 2 | 3.96 | .049† |
| Amygdala | R | 20 | -8 | -16 | 3.80 | .017† |
| | L | -30 | -2 | -24 | 3.63 | .030† |
| *Nocebo > Placebo* | | | | | | |
| Thalamus | R | 6 | -6 | 2 | 3.90 | .048† |

*Note.* Coordinates are in MNI space. SMG = Supramarginal Gyrus. DLPFC = Dorsolateral Prefrontal Cortex. STG = Superior Temporal Gyrus. vmPFC = ventromedial prefrontal cortex. † small-volume corrected.

**Supplementary Table 5**

*Peak values of effects found in the combined EEG-fMRI analysis*

| | | | Peak | | | |
|---|---|---|---|---|---|---|
| **Region** | **Hemi.** | **Direction** | **Elec.** | **Freq.** | **Time** | ***t(40)*** |
| Anterior Insula | L | negative | F8 | 16 Hz | -3.0 s | -5.13 |
| Anterior Cingulate Cortex | L | positive | PO3 | 128 Hz | -2.1 s | 5.34 |
| | | positive | TP10 | 128 Hz | 0.2 s | 4.89 |
| | R | positive | FT7 | 76.11 Hz | -0.1 s | 5.35 |
| DLPFC | L | negative | F5 | 26.89 Hz | -5.2 s | -5.47 |

**Note.** Elec = Electrode. Freq = Frequency. DLPFC = Dorsolateral Prefrontal Cortex. All peak *t*-values were significant at $p < .001$.
